# Oat chromosome and genome evolution defined by widespread terminal intergenomic translocations in polyploids

**DOI:** 10.1101/2022.08.23.504991

**Authors:** Paulina Tomaszewska, Trude Schwarzacher, Pat (J.S) Heslop-Harrison

## Abstract

Structural chromosome rearrangements involving translocations, fusions and fissions lead to evolutionary variation between species and potentially reproductive isolation and variation in gene expression. While the wheats (Triticeae, Poaceae) and oats (Aveneae) all maintain a basic chromosome number of *x*=7, genomes of oats show frequent intergenomic translocations, in contrast to wheats where these translocations are relatively rare. We aimed to show genome structural diversity and genome relationships in tetraploid, hexaploid and octoploid *Avena* species and amphiploids, establishing patterns of intergenomic translocations across different oat taxa using fluorescence *in situ* hybridization (FISH) with four well-characterized repetitive DNA sequences: pAs120, AF226603, Ast-R171 and Ast-T116. In *A. agadiriana* (2*n*=4*x*=28), the selected probes hybridized to all chromosomes indicating that this species originated from one (autotetraploid) or closely related ancestors with the same genomes. Hexaploid amphiploids were confirmed as having the genomic composition AACCDD, while octoploid amphiploids showed three different genome compositions: AACCCCDD, AAAACCDD or AABBCCDD. The A, B, C, and D genomes of oats differ significantly in their involvement in non-centromeric, intercalary translocations. There was a predominance of distal intergenomic translocations from the C-into the D-genome chromosomes. Translocations from A- to C-, or D- to C-genome chromosomes were less frequent, proving that at least some of the translocations in oat polyploids are non-reciprocal. Rare translocations from A- to D-, D- to A- and C- to B-genome chromosomes were also visualized. The fundamental research has implications for exploiting genomic biodiversity in oat breeding to through introgression from wild species potentially with contrasting chromosomal structures and hence deleterious segmental duplications or large deletions in amphiploid parental lines.

## Introduction

Polyploidy and whole genome duplication have been recognized as major evolutionary processes in plants (Soltis et al., 2015; Alix et al., 2017; Zwaenepoel and Van de Peer, 2019; Heslop-Harrison et al., 2022). While all plants are known to have whole genome duplications within their ancestry, one or more post cretaceous-tertiary (K-T) polyploidy events have been found in about half of species, including crops and wild plants. Genes that have been duplicated during the polyploidization process may retain or change their original function and can be mutationally or epigenetically silenced. In new polyploids, many evolutionary processes occur above the organizational level of duplicated genes (Adams and Wendel, 2005). These include elimination of whole chromosomes or even whole genomes (*Hordeum*: Gernand et al., 2005; *Nicotiana:* Patel et al., 2011) as well as intra- and inter-genomic chromosome translocations (Badaeva et al., 2007; Tomaszewska, 2021; Tomaszewska and Kosina, 2021). Changes occurring in polyploid nuclei may be associated with transposon activity (McClintock, 1984; Comai et al., 2003) and lead to the stabilization of polyploid genomes, thus increasing vitality and fertility, and extending the adaptive potential of such plants (Alix et al., 2017).

Chromosome translocations are important for plant evolution (Martin et al., 2020; Wang et al., 2021; Biswas et al., 2022; Liu et al., 2022), and occur in diploids, and after hybrid or amphiploid formation; they lead to exchange of chromosome segments within and between the ancestral genomes. Comparative genetic analysis in the grass family, including major crops, showed that particular taxonomic units are characterized by different chromosome translocations in terms of their position on chromosomes, reciprocity, and number of breaks involved (Bardsley et al. 1999). Kubaláková et al. (2003) described various intragenomic reciprocal translocations in rye. Søgaard & von Wettstein-Knowles (1987) found intragenomic translocations located at or near the centromere, or interstitially as the most common in the barley genome. Badaeva et al. (2007) showed that single intragenomic translocations between the B-genome chromosomes were the most frequent in wheat species. A detailed analysis of 373 Chinese wheat varieties has shown 14 different structural chromosome rearrangements including single- and reciprocal-translocations (Huang et al., 2018). Only two types of intergenomic translocations between A and B-genome chromosomes were observed regularly in 4*x* and 6*x* wheats. Translocations between D- and A- or B-genome (including 1Ds chromosomes) were rare, and most translocation breakpoints were at or near the centromere, rarely interstitially. These data together with the analysis of pericentric inversions conducted by Qi et al. (2006) give insight into the evolutionary dynamics of pericentromeric regions in the Triticeae tribe. The outcomes of genome evolution are different in the sister tribe of Aveneae, also involving an invariant basic chromosome number of *x*=7 and various polyploids: the oat lineage originated before wheat, with the C genome appearing earlier than the A or D genomes, suggesting a longer evolution time for oat hexaploids than for wheat (Fu, 2018; Peng et al., 2022), certainly with the major hexaploid crop species *Triticum aestivum*.

The genus *Avena* comprises about thirty diploid, tetraploid and hexaploid species with a basic chromosome number *x*=7 (Baum, 1977). The genomes of oat species were classified into A, B, C and D genome groups and further subdivisions (Rajhathy and Thomas, 1974; designated by a subscript e.g., A_l_, A_s_, C_p_ or C_v_). Diploid species are characterized by the presence of A or C genomes; the B genome is found only in some tetraploids. The D genome is present in all hexaploids (Badaeva et al., 2005) and some tetraploids (Yan et al., 2016, 2021; Tomaszewska and Kosina, 2021). The literature on the number and localization of intergenomic chromosome translocations in oats mainly uses *in situ* hybridization with total genomic DNA or some abundant repetitive DNA sequences (Chen and Armstrong, 1994; Jellen et al., 1994; Leggett et al., 1994; Linares et al., 2000; Tomaszewska and Kosina, 2021). Many distal (terminal) and several interstitial (subterminal) intergenomic translocations were observed, and only a few of them were reciprocal. Although the detection of translocation between A/D and C or A and B genomes with the use of genomic *in situ* hybridization, or genome-specific repetitive DNA probes, is straightforward, it is much more difficult to determine the translocations between A and D genomes due to the lack of sequence differentiation (Katsiotis et al., 2000; Luo et al., 2014; Liu et al., 2019; see wheat results from Huang et al., 2018). Linares et al. (1998) distinguished A and D genomes with the use of satellite sequences specific for the A genome, and Liu et al. (2019) developed probes specific for the D-genome. However, the translocations between A and D genomes have only been observed in oat endosperm (Tomaszewska and Kosina, 2021).

The combination of molecular cytogenetic and genomic or bioinformatic methods are promising to identify the processes occurring when genomes come together in polyploids (Heslop-Harrison and Schwarzacher, 2011; Tomaszewska et al., 2022). We can then address features of polyploidy that may be related to evolutionary adaptation (Barker et al., 2016) and test the hypotheses that chromosomal rearrangements, including intergenomic translocations, occur early in polyploid evolution. For this purpose, it is worth using artificial hybrids and amphiploids to track changes in genomes shortly after hybridization and duplication, and in further progenies (Tomaszewska and Kosina, 2018, 2021). Understanding the coexistence of genomes in polyploid nuclei and consequences of genome rearrangement in polyploids is important to answer how non-cultivated diploids can be used for germplasm enhancement. Following either natural selection, speciation, or laboratory crossing, diploid genomes may be stabilized in a polyploid, or in backcrosses regularly used to generate breeding lines including wild species (Newell et al., 2012) so lines made for research may be more widely valuable. Translocations and other rearrangements may be useful to ensure desirable gene combinations are retained during breeding or represent a challenge because they restrict recombination.

Here we aim to recognize genomes in oat diploids and identify sequence characteristics of genomes with respect to the repetitive DNA composition, to show genome-relationships in tetra- and hexaploid *Avena* species and establish genome composition of their synthetic hybrids and amphiploids (6*x* and 8*x*) using fluorescence *in situ* hybridization (FISH) with well-characterized repetitive DNA sequences. Then we aim to establish patterns of intergenomic translocations across different taxa and relate it to the evolutionary divergence of oat genomes.

## Materials and methods

### Plant materials

Species, hybrids and amphiploids of oats used in the study are listed in **Table 1**. Accessions were kindly provided by international germplasm collections. Some amphiploids and hybrids studied here were developed in United Kingdom, and also by Nishiyama in Japan mostly in the 1960-1980s (Nishiyama, 1962; Nishiyama et al., 1963; Nishiyama and Yabuno, 1979). The botanical nomenclature of studied oat species was applied according to http://www.theplantlist.org/, accessed on 16 August 2022.

**Table 1.**
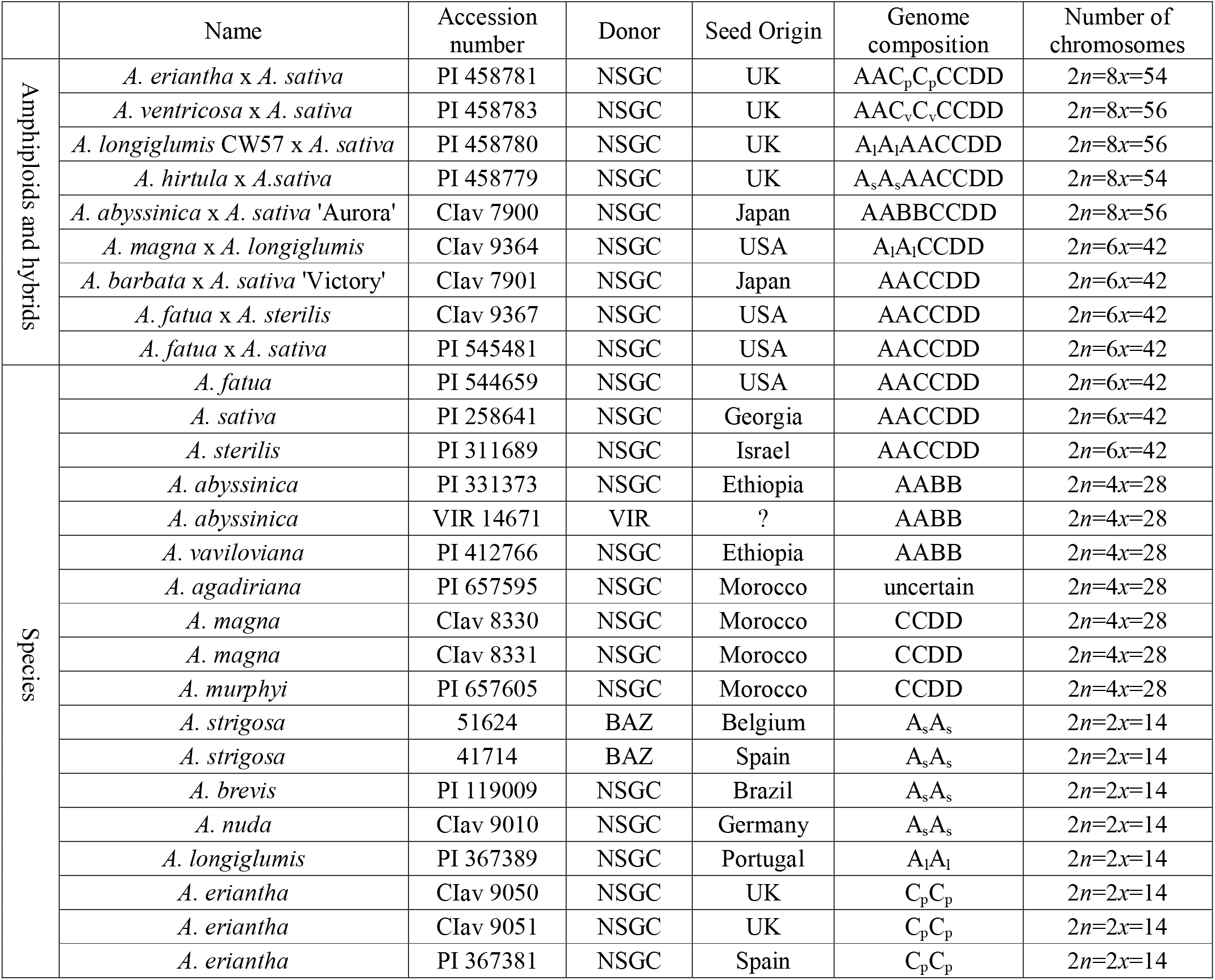
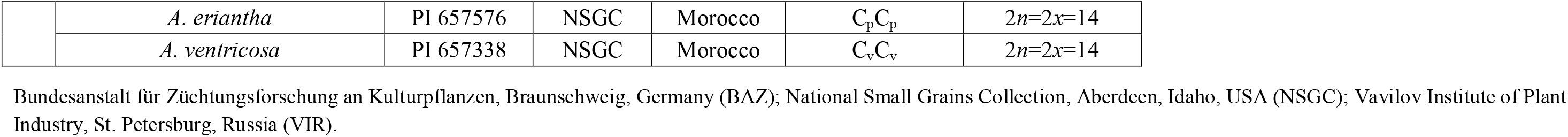
List of species, hybrids and amphiploids of oats used in the study.

### Chromosome preparation

Oat chromosomes were prepared according to the protocol described by Schwarzacher and Heslop-Harrison (2000). The root tips were treated with ice-cold water for 24 h to accumulate metaphases, and fixed in 96% ethanol: glacial acetic acid (3:1) for 48 h. Fixed root-tips were then washed in enzyme buffer (10mM citric acid/sodium citrate) for 15 min, and digested in an enzyme solution composed of 20U/ml cellulase (Sigma C1184), 10U/ml Onozuka RS cellulase (RPI C32400), and 20U/ml pectinase (Sigma P4716 from *Aspergillus niger*; solution in 40% glycerol) for 60 min at 37°C. Root tips were then squashed in 60% acetic acid. Cover slips were removed after freezing with dry ice. Slides were air-dried and used for *in situ* hybridization.

### Probes used for *in situ* hybridization

Four different probes were selected for fluorescence *in situ* hybridization to distinguish genomes in polyploids and recognize major intergenomic translocations:

1. A genome-specific pAs120 (Linares et al., 1998)
2. C genome-specific AF226063 (Ananiev et al., 2002; Liu et al., 2019)
3. D genome-specific Ast_R171 (Liu et al., 2019)
4. D genome-specific Ast_T116 (Liu et al., 2019)

Conserved regions were amplified in a standard Polymerase Chain Reaction (PCR) using genome-specific primers (Linares et al., 1998; Liu et al., 2019) synthesized commercially (Sigma-Aldrich). These probes were labelled with digoxigenin-11-dUTP, biotin-16-dUTP or tetramethyl-rhodamine-5-dUTP (Roche) using BioPrime Array CGH and purified using BioPrime Purification Module (Invitrogen). The 45bp AF226063 oligonucleotide probe (Liu et al. 2019) was synthesized commercially (Sigma-Aldrich) with TET fluorescent dye attached to oligonucleotides at the 5’-end.

### Fluorescence *in situ* hybridization procedure

FISH was performed as described by Schwarzacher and Heslop-Harrison (2000) and Tomaszewska and Kosina (2021) with minor modifications. The hybridization mixture consisted of 50% deionised formamide, 10% dextran sulphate, 1% sodium dodecyl sulphate (SDS), 2X SSC (saline sodium citrate buffer), amplified and labelled probe(s) (2 ng μL^-1^ each), and 200 ng μL^-1^ salmon sperm DNA was predenatured for 10 min at 75°C and stabilized on ice for 10 min. The 45bp AF226063 oligonucleotide probe was not predenatured and was added to the hybridization mixture containing amplified probes shortly after the predenaturation step. The hybridization mixture and chromosomes were then denatured together in a hybridization oven for 7 min at 75°C. Hybridization was performed at 37°C overnight. Slides were washed at 42°C in 2X SSC for 2 min, in 0.1X SSC for 10 min, and 2X SSC for 20 min. Hybridization signals of probes labelled with digoxigenin-11-dUTP and biotin-16-dUTP were detected with antidigoxigenin conjugated to fluorescein isothiocyanate (FITC; Roche Diagnostics) and streptavidin conjugated to Alexa 594 or Alexa 647 (Life Technologies-Molecular Probes), respectively. Air-dried slides were counterstained with DAPI (4’,6-diamidino-2-phenylindole, 2μg mL) in antifade solution (AF1, Citifluor).

### Microscopy and image capture

The slides were examined with a Nikon Eclipse 80i epifluorescence microscope (Nikon, Tokyo, Japan). Images were taken using a DS-QiMc monochromatic camera and NIS-Elements v.2.34 software. Karyotypes were prepared using IdeoKar 1.3 (Mirzaghaderi and Marzangi, 2015) and Adobe Photoshop.

## Results

### Chromosome numbers

The number of chromosomes of studied accessions is shown in **Table 1**. For FISH analysis, we used accessions with the number of chromosomes typical for a given species and thus omitted *A. wiestii* PI 299112 and *A. barbata* PI 337795, each having 42 chromosomes instead of the reported 14 and 28, respectively, suggesting uncontrolled crossing in breeding or involvement of 2*n* gametes in stocks. All species, hybrids and amphiploids of oat studied here were euploid (**Table 1**), except *A. eriantha* × *A. sativa* and *A. hirtula* × *A. sativa* where most metaphases had 2*n*=8*x*-2=54. Ploidy levels of hybrids and amphiploids were re-examined and compared with genebank databases. Ploidy level of *A. barbata* × *A. sativa* ‘Victory’ CIav 7901, previously recognized as octoploid, had to be revised to hexaploid.

### FISH-based reference karyotypes of diploid oats

Two different probes, pAs120 (Linares et al., 1998) and AF226063 (Ananiev et al., 2002; Liu et al., 2019) repeats, were used for *in situ* hybridization on metaphases of diploid oat species (**Table 1**). The karyotypes were prepared, and the patterns of signals of the two genome-specific probes were determined (**Fig. 1**). These patterns were helpful in establishing the genomic composition of polyploids and recognizing intergenomic translocations in tetra-, hexa- and octo-ploids. Probe pAs120 has been tested on chromosomes of different diploids having A_l_ (*A. longiglumis*) and A_s_ genomes (*A. brevis, A. nuda, A. strigosa*). All studied species showed dispersed signals along chromosomes, except for telomeres and secondary constrictions. Probe pAs120 was not useful for distinguishing between A_l_ and A_s_ genomes. Probe AF226063 has been used on chromosomes of different accessions of diploid *A. eriantha* (C_p_ genome) and *A. ventricosa* (C_v_ genome). Some chromosomes showed specific probe signals in the form of bands being good chromosome markers. Probe AF226063 was not useful for distinguishing between C_p_ and C_v_ genomes.

**Fig. 1.**
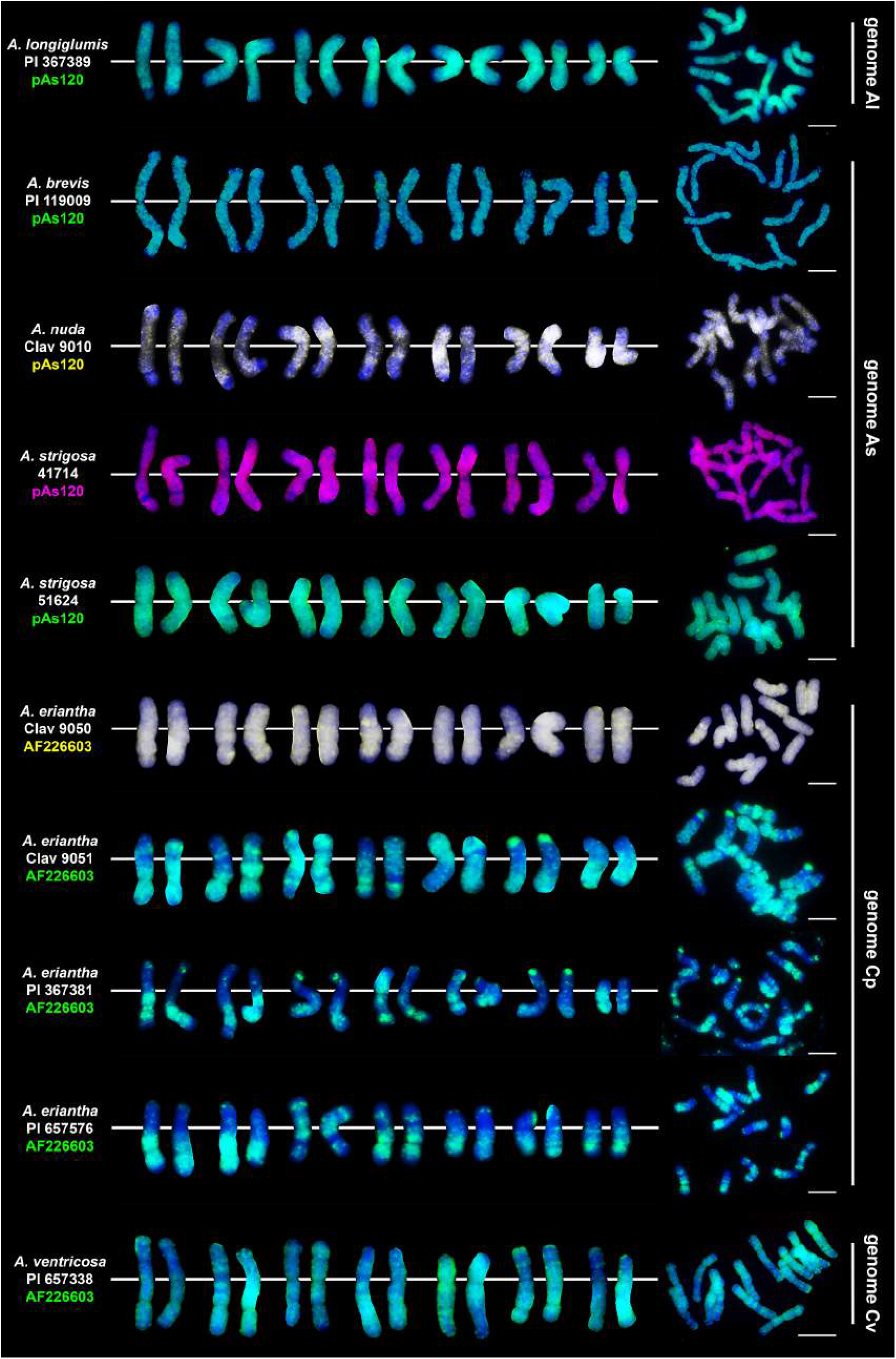
Chromosomal location of highly repetitive DNA motifs in diploid *Avena* species having genome constitution of A_l_, A_s_, C_p_ and C_v_. Scale bar = 10μm

### Chromosomal location of highly repetitive DNA motifs in tetraploid and hexaploid oat species

We studied three species belonging to the AABB *Avena* group: *A. abyssinica, A. vaviloviana* and *A. agadiriana*. Genome composition of *A. abyssinica* and *A. vaviloviana* was confirmed using pAs120 probe, which painted 14 chromosomes out of 28, designating the A genome (**Fig. 2**). Chromosomes that were not painted by pAs120 were considered genome B. Hybridization of Ast-T116 probe to chromosomes of *A. agadiriana* gave strong dispersed signals along all of the 28 chromosomes. Another probe pAs120 gave weak dispersed signals along 28 chromosomes, hence the genomic composition of this species remains ambiguous. Genome composition of two tetraploid species previously classified into AACC *Avena* group, *A. magna* (syn. *A. maroccana*) and *A. murphyi*, was established using two different probes (**Fig. 3**). In both species, AF226603 painted 14 chromosomes out of 28 while Ast-T116 showed dispersed signals along the remaining 14 chromosomes, proving that the genome composition of these species was CCDD. All hexaploid species studied here, including *A. fatua, A. sterilis* and *A. sativa,* had AACCDD genome composition, and this was confirmed using three different genome-specific probes: pAs120, AF226603 and Ast-T116 (**Fig. 4**).

**Fig. 2.**
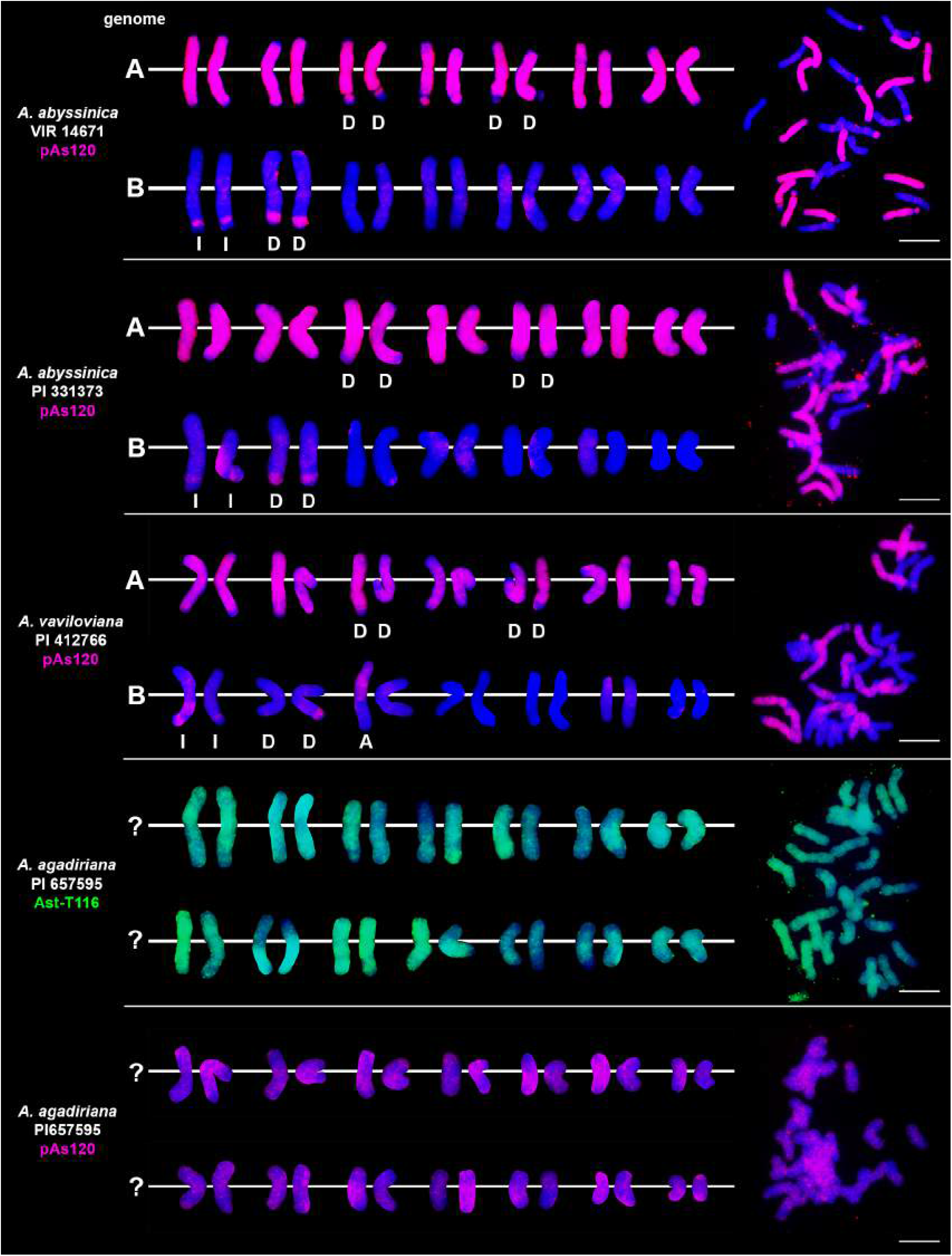
Genome composition and intergenomic translocations pattern in tetraploid oat species belonging to AABB *Avena* group. D-distal translocation, I-interstitial translocation, A - translocation of whole arm. Scale bar = 10μm

**Fig. 3.**
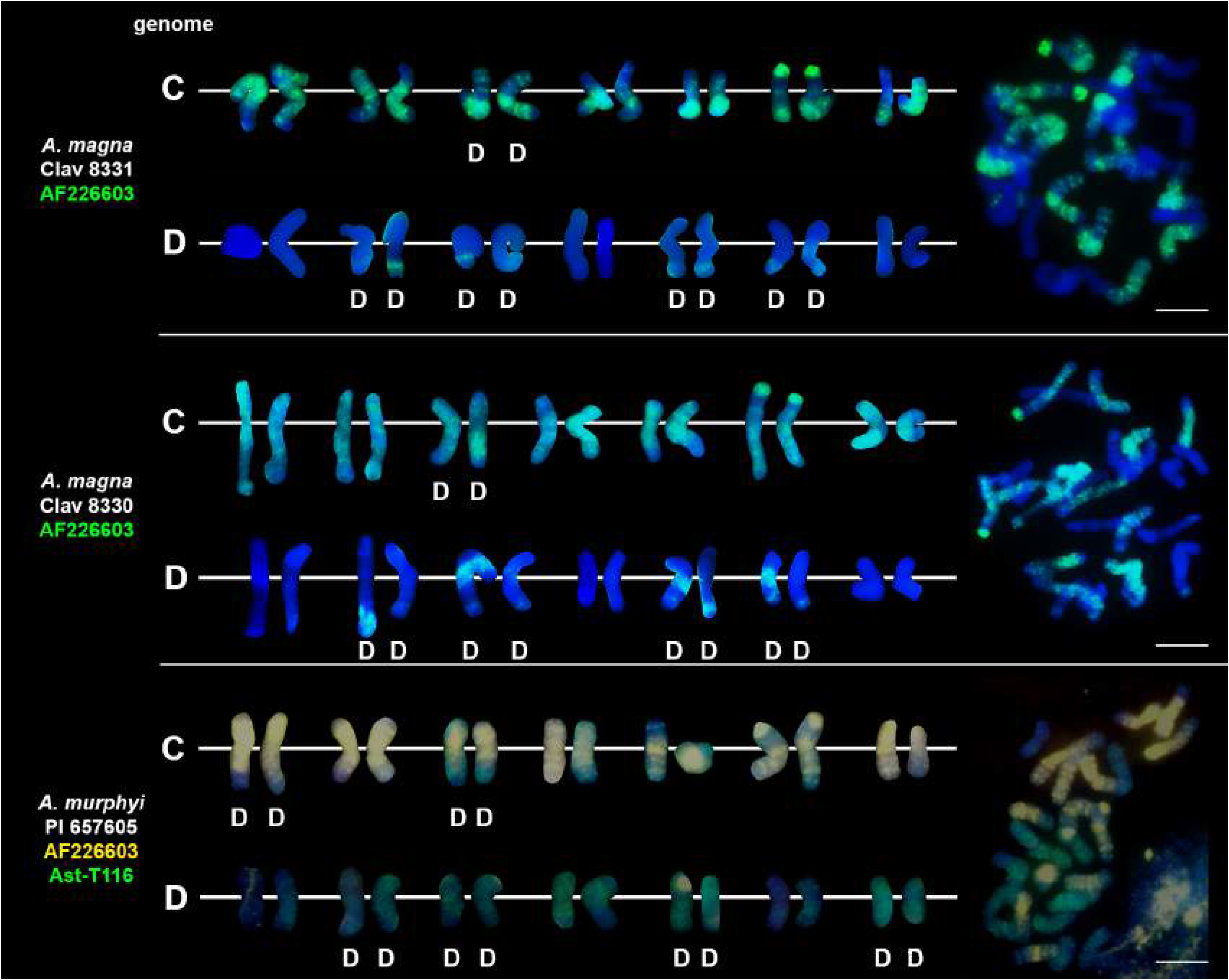
Genome composition and intergenomic translocations pattern in tetraploid oat species belonging to CCDD *Avena* group. D-distal translocation. Scale bar = 10μm

**Fig. 4.**
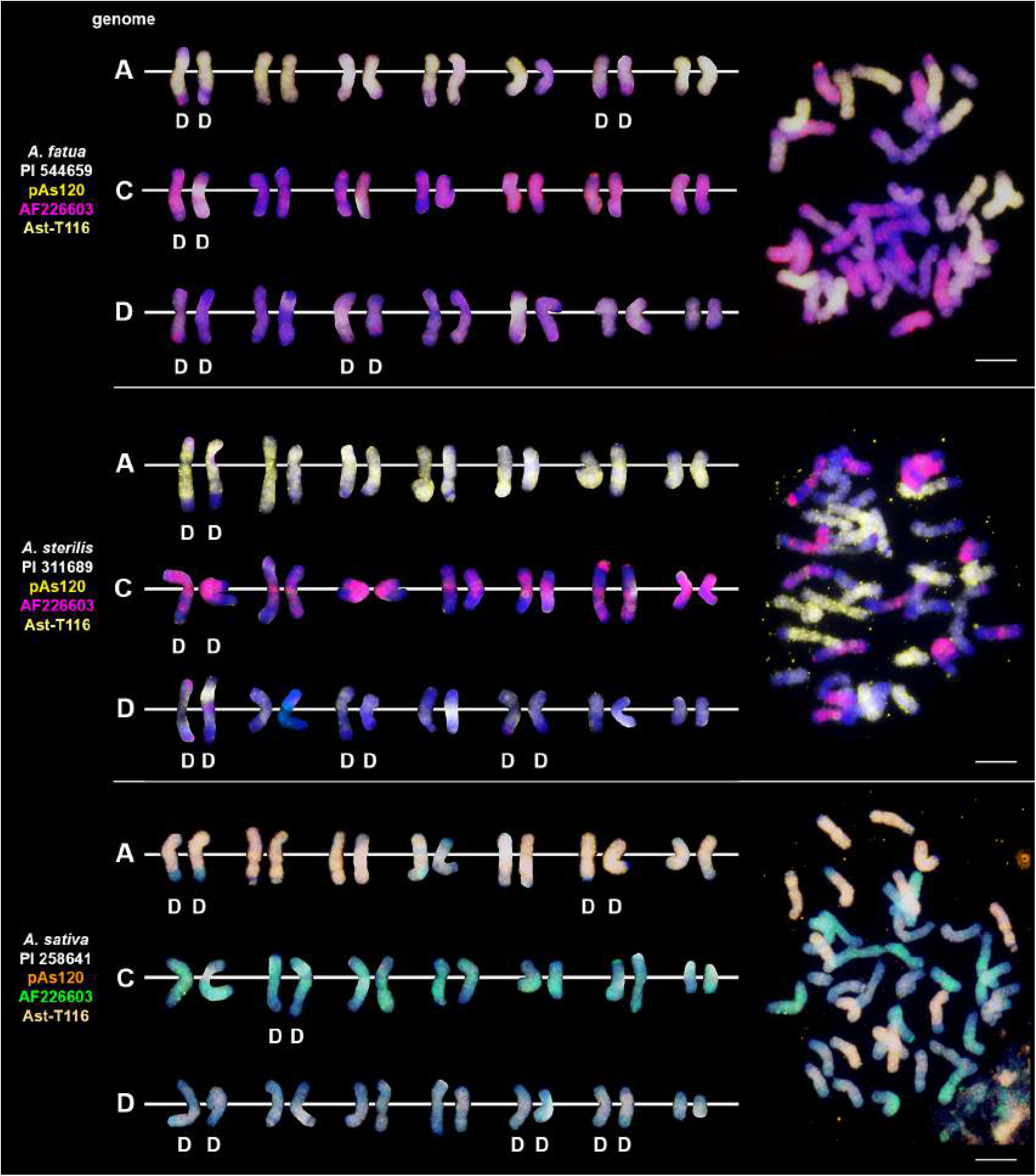
The diversity of intergenomic chromosome translocations across different hexaploid oat species. D-distal translocation. Scale bar = 10μm

### Chromosomal organization of specific repeats in artificial hybrids and amphiploids of oats

Probes pAs120, AF226603, Ast-R171 and Ast-T116 were used to establish genome composition of different artificial hybrids and amphiploids of oats. The probes produced multiple signals that were evenly distributed along 14 chromosomes each, enabling identification of A, C and D genomes. Chromosomes that were not painted by any of these probes were considered genome B. Each of the analyzed hexaploid hybrids and amphiploids had the genomic composition of AACCDD (**Fig. 5**), as did hexaploid oat species (**Fig. 4**). In the amphiploid *A. magna* × *A. longiglumis*, we were able to more accurately determine the genomic composition as A_l_A_l_CCDD by looking at the genomes of the parental species. We recognized 3 types of octoploid amphiploids, each having different genome composition: AACCCCDD, AAAACCDD or AABBCCDD. *A. eriantha* × *A. sativa* and *A. ventricosa* × *A. sativa* belonged to the first group having AAC_p_C_p_CCDD and AAC_v_C_v_CCDD genomes, respectively (**Fig. 6**). No letter in the subscript was assigned to the second C genome of these amphiploids because this genome originated from *A. sativa* where the ancestors of this species have not been thoroughly investigated. *A. longiglumis* CW57 × *A. sativa* and *A. hirtula* × *A.sativa* belonged to the second group of amphiploids having A_l_A_l_AACCDD and A_s_A_s_AACCDD genomes, respectively (**Fig. 7**). No letter in the subscript was assigned to the second A genome originated from *A. sativa*. The most likely genomic composition of *A. abyssinica* × *A. sativa* ‘Aurora’ is AABBCCDD (**Fig. 8**). One of the genomes of this amphiploid has not been labelled with any probe and is believed to belong to genome B.

**Fig. 5.**
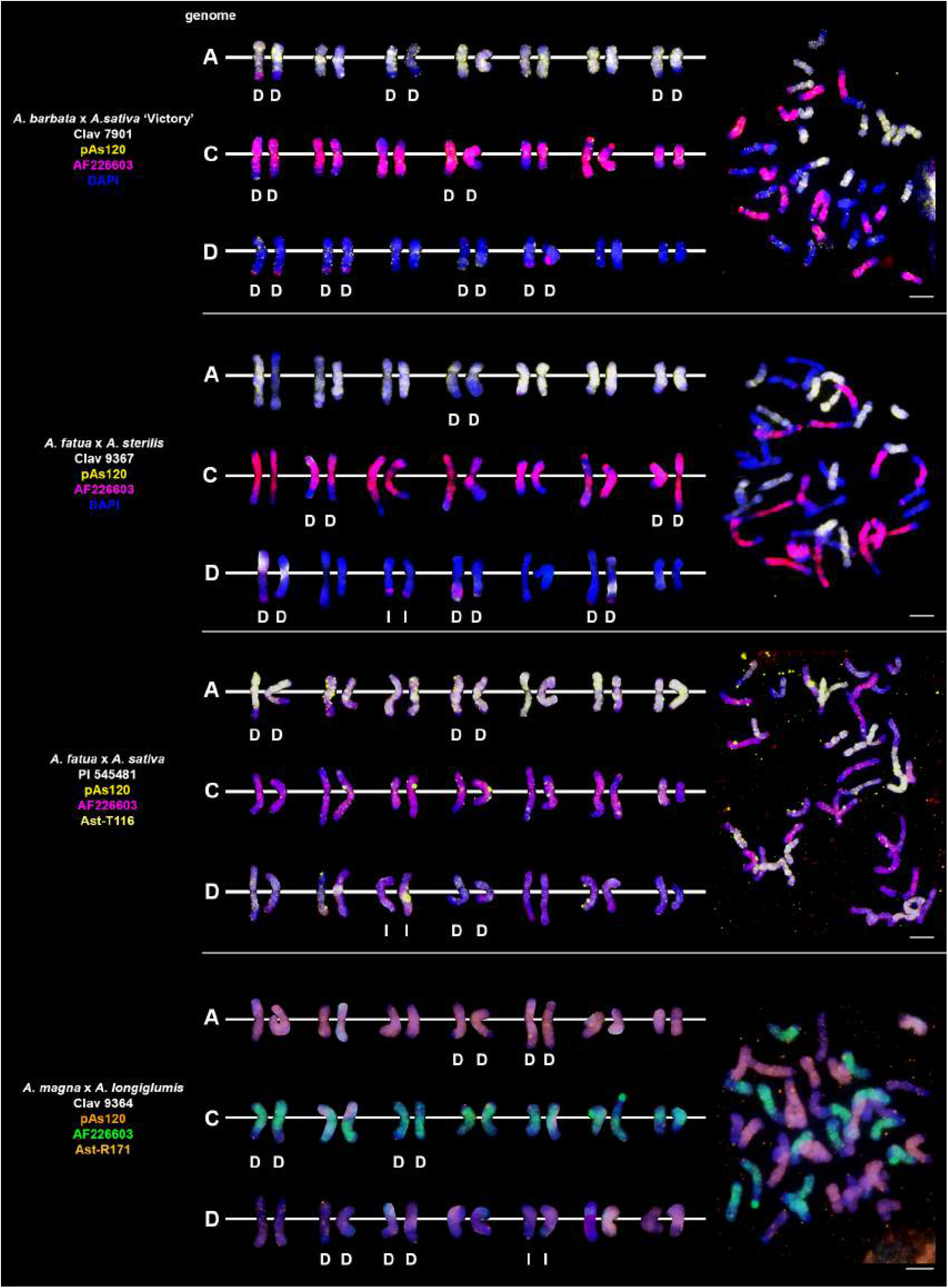
Chromosomal location of repetitive DNA motifs enabling identification of AACCDD genomes and intergenomic translocations in different hexaploid hybrids and amphiploids of oats. D-distal translocation, I-interstitial translocation. Scale bar = 10μm

**Fig. 6.**
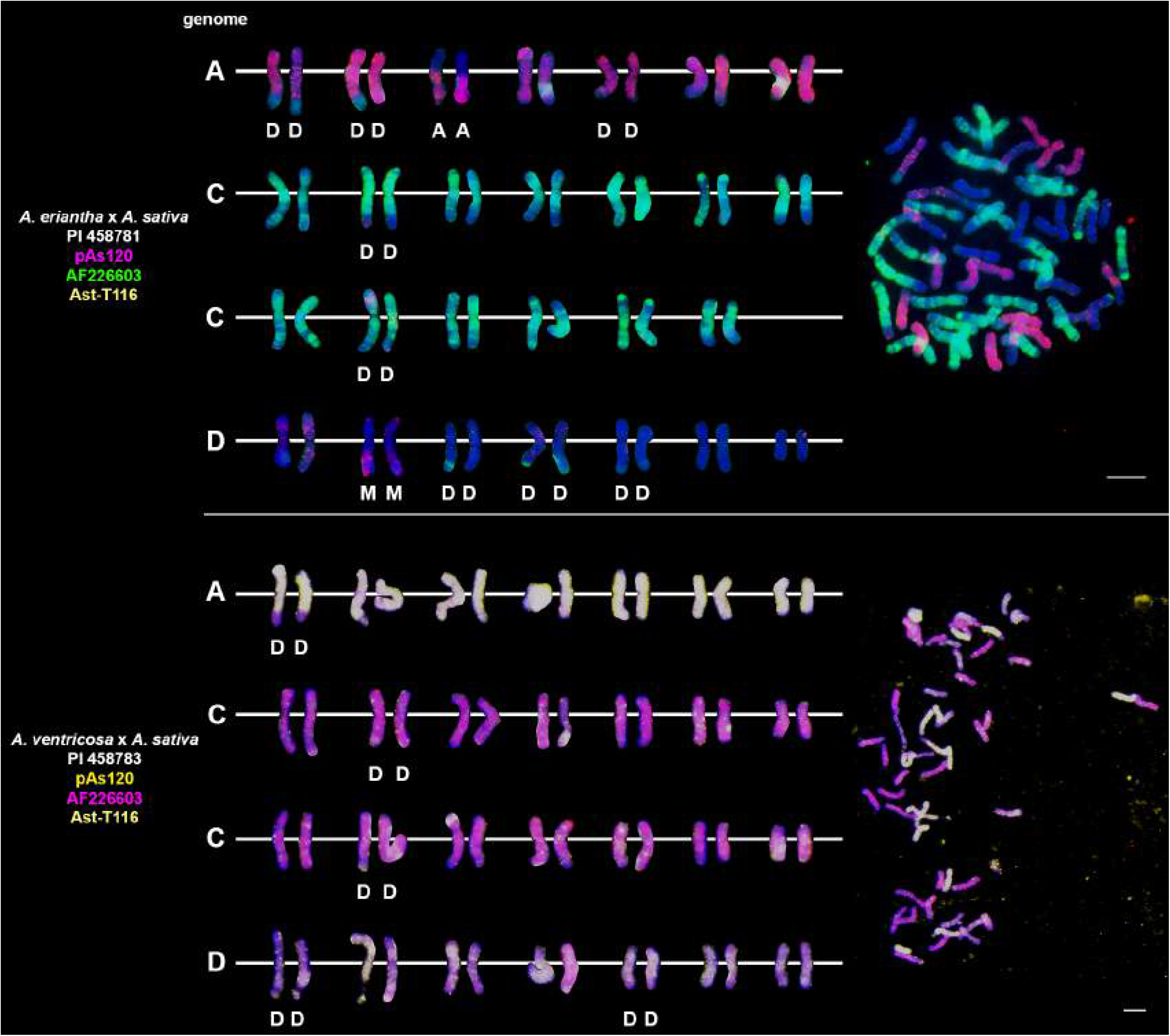
Fluorescence *in situ* hybridization with repetitive DNA probes to chromosomes of amphiploids formed by crossing diploid species having CC genomes with *A. sativa*. Intergenomic translocations were visualized. D-distal translocation, I-interstitial translocation, M - multiple translocation or insertion. Scale bar = 10μm

**Fig. 7.**
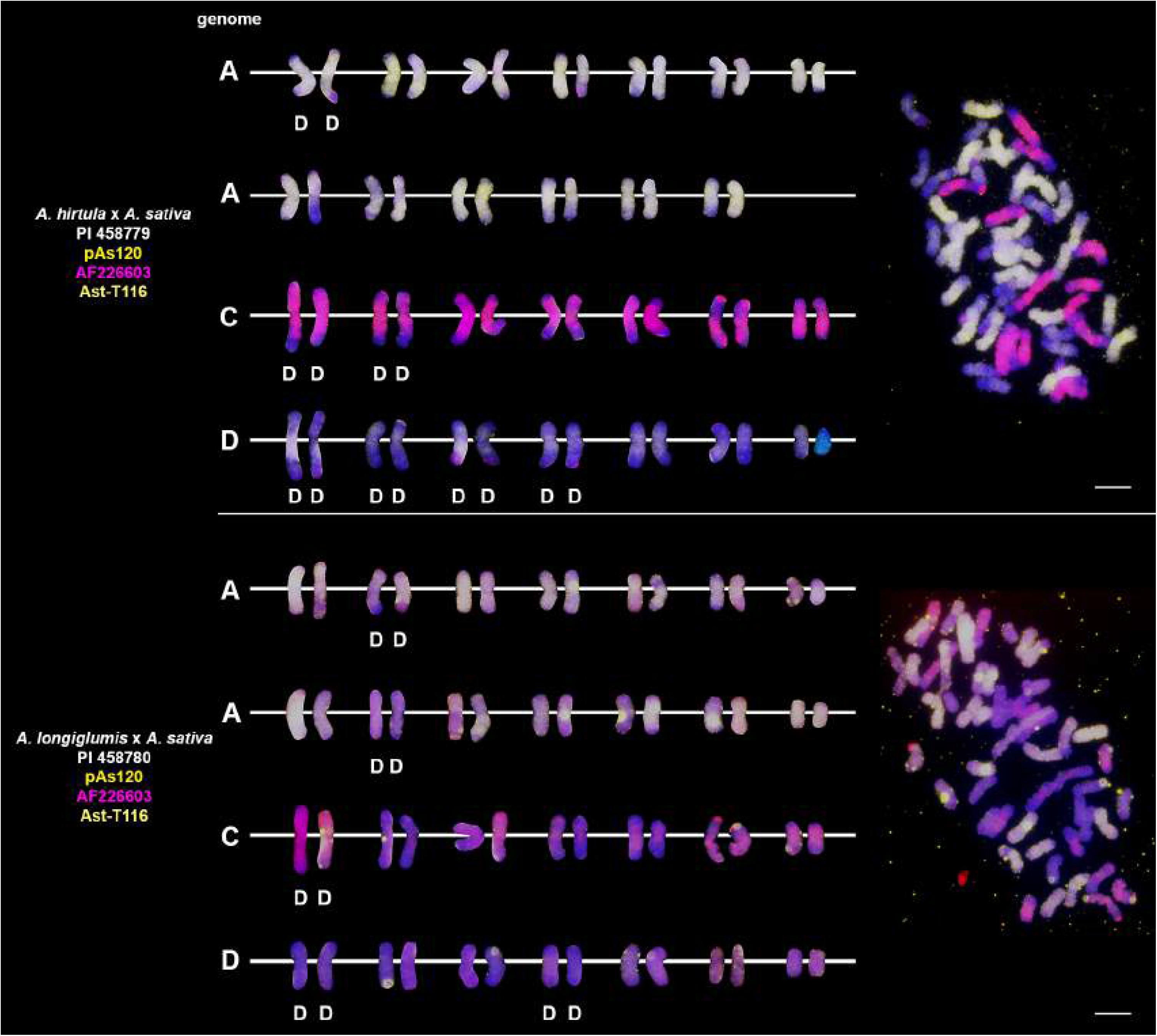
Fluorescence *in situ* hybridization with repetitive DNA probes to chromosomes of amphiploids formed by crossing diploid species having AA genomes with *A. sativa*. Intergenomic translocations were visualized. D-distal translocation. Scale bar = 10μm

**Fig. 8.**
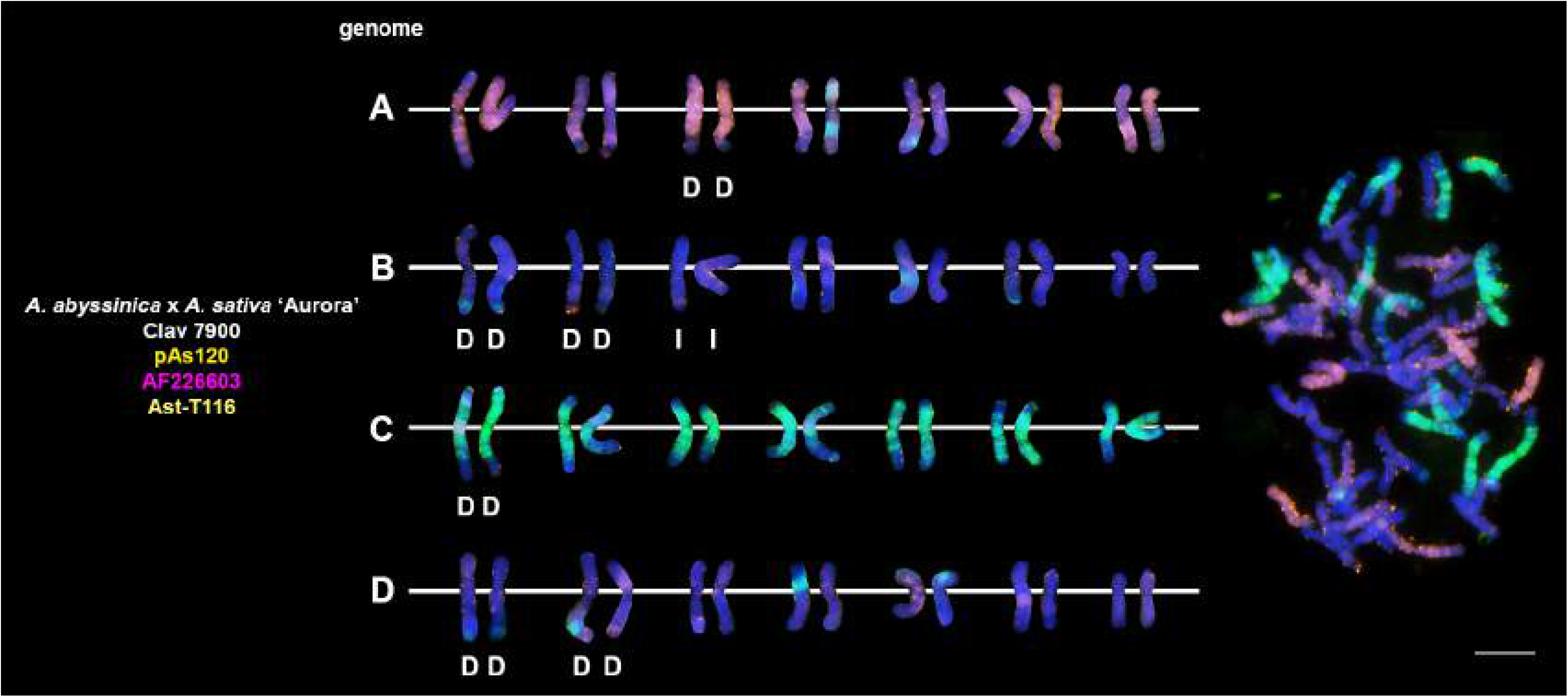
Fluorescence *in situ* hybridization with repetitive DNA probes to chromosomes of an amphiploid formed by crossing tetraploid *A. abyssinica* with *A. sativa*. Intergenomic translocations were visualized. D-distal translocation, I-interstitial translocation. Scale bar = 10μm

### Visualization of intergenomic translocations using genome-specific repeats

Fluorescent *in situ* hybridization with the use of repetitive DNA motifs to chromosomes of different polyploids (4*x,* 6*x*, 8*x*) enabled identification of intergenomic translocations (**Figs. 2–8**). Using probes specific to genomes A, C and D, we were able to discriminate C→A, C→D and A→C, D→C translocations in hexaploids (**Figs. 4, 5**) and octoploids (**Figs. 6–8**). Rare A→D, D→A and C→B translocations were also visualized. The translocations involving B and D genomes were not detected in *A. abyssinica* × *A. sativa* ‘Aurora’ octoploid (**Fig. 8**). Most of the translocation events were at the distal position of long chromosome arms (terminal or subterminal), rarely interstitially (**Figs. 2–8**). Some of the distal translocations involved SAT chromosomes, as observed in *A. abyssinica* and *A. vaviloviana* (**Fig. 2**). In some of the accessions examined, we detected whole chromosome arm translocations, i.e. *A. vaviloviana* (**Fig. 2**; one chromosome from a pair) or *A. eriantha* × *A. sativa* (**Fig. 6**; two chromosomes from a pair). In the latter, we additionally revealed a pair of D genome chromosomes having red C genome signals at the ends of both arms. It suggests multiple distal translocations or insertion, presumably involving chromosome 3 from genome A showing large whole arm translocation (**Fig. 6**).

### Pattern of intergenomic translocations across different oat taxa

Our detailed studies of genome relationships in polyploid species and interspecific amphiploids and hybrids of oats, including both FISH (**Figs. 2–8**, **Table 2**) and some literature data (**Table 2**) analyses, indicated that the chromosomes of A, B, C and D genomes differ significantly in their involvement in translocations. There is a predominance of distal intergenomic translocations from the C-into the D-genome chromosomes in CCDD-tetraploid and AACCDD-hexaploid species. A→C or D→C translocations are less frequent, proving that at least some of the translocations in oat polyploids are non-reciprocal. The number of A→B and B→A translocations in tetraploid *A. abyssinica* was the same, but their position on the chromosomes (distal *versus* interstitial) indicated their non-reciprocity.

**Table 2.**
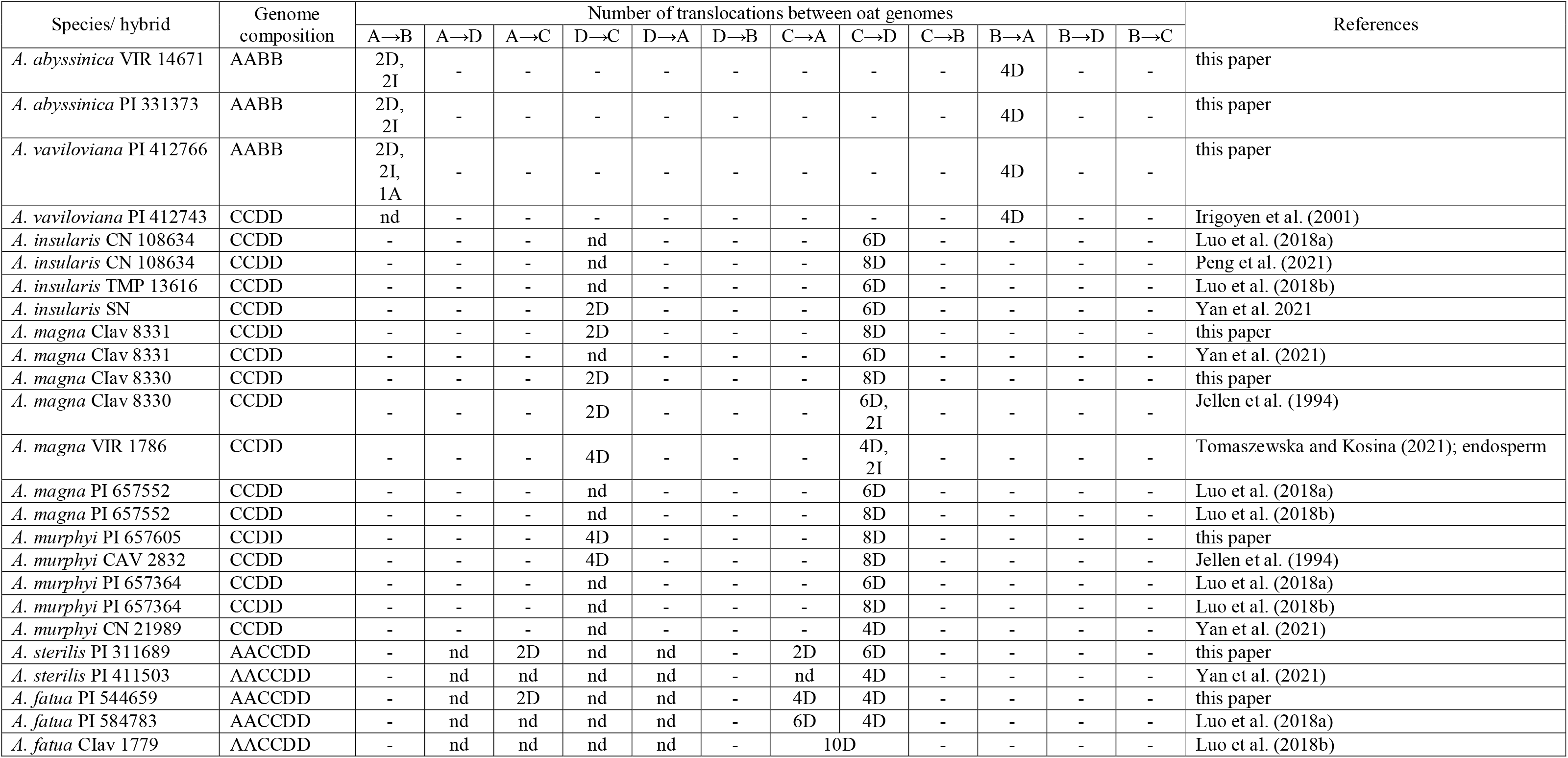
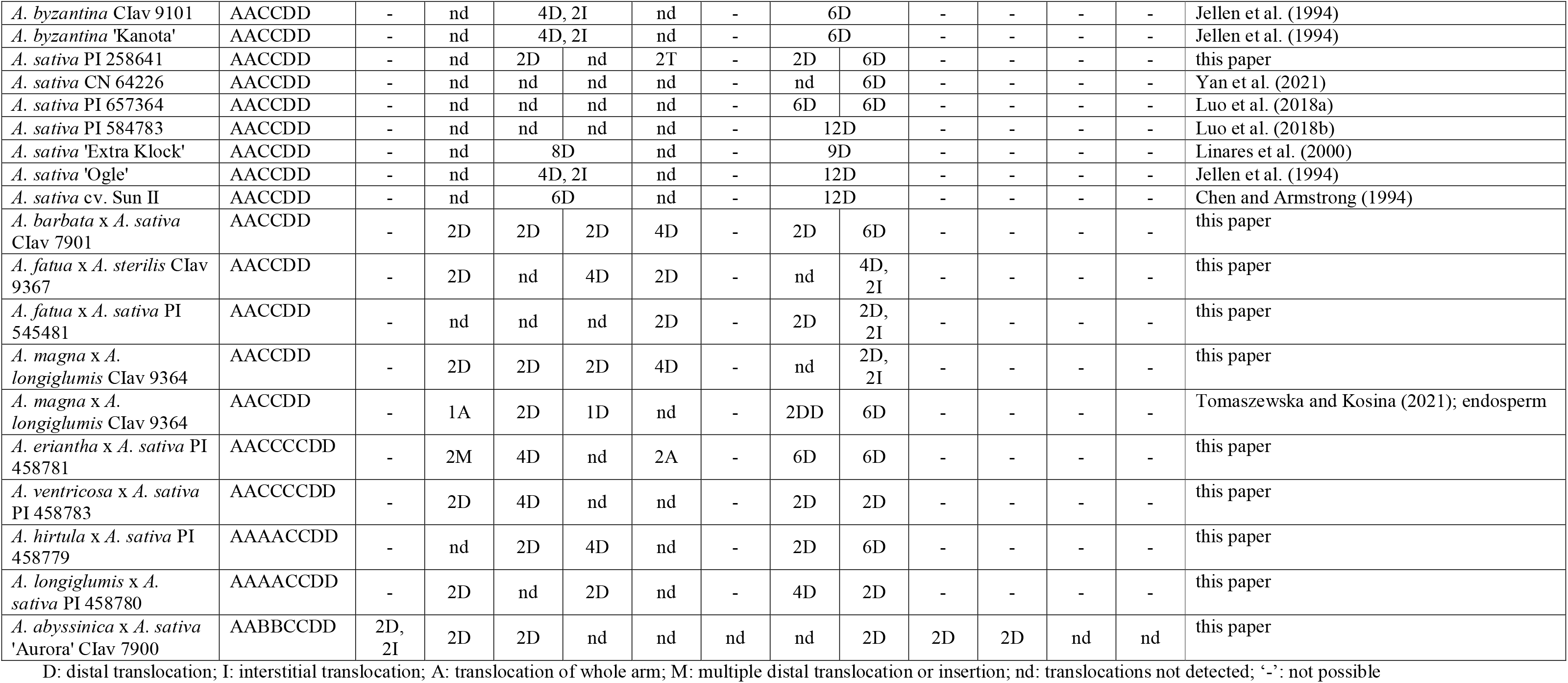
Pattern of intergenomic translocations across different oat taxa.

Comparative analysis of species and artificial hybrids/amphiploids having genomic composition of AACCDD (**Figs. 4, 5**; **Table 2**) showed that these two groups differ in the number and position of translocations. Multitude of translocations covering all genomes in hybrid octoploids (**Figs. 6–8**), and probably other extensive changes including large insertions (*A. eriantha* × *A. sativa,* **Fig. 6**) have occurred soon after amphiploid formation.

Our analysis across different oat taxa (**Figs. 2–8**; **Table 2**) showed that some species, such as *A. abyssinica* and *A. vaviloviana* (**Fig. 2**), share some of the intergenomic translocations (most likely they are identical-by-descent, but re-iteration of same events in different lineages cannot be ruled out). The translocation of the entire arm of a single chromosome observed in *A. vaviloviana* may be species-specific or specific for this particular accession. Both our karyotype analysis (**Fig. 3**) and the data obtained by different authors (**Table 2**) indicated that some translocations between the C and D genomes in hexaploid oat species (**Fig. 4**) involved the same chromosomes as in tetraploid CCDD species (**Fig. 3**). The translocation pattern seen in amphiploids created by crossing different maternal diploids with paternal *A. sativa* indicated that some translocations were transferred from *A. sativa* to the hybrid generation, and new translocations were also appeared (**Figs. 5–7**).

## Discussion

Repetitive DNA elements are rapidly evolving major components of plant genomes, thus becoming important tools for studying the large-scale organization and evolution of plant genomes (Heslop-Harrison and Schwarzacher, 2011; Mehrotra and Goyal, 2014; Biscotti et al., 2015). Here, by using identified repetitive sequences that were unique to the A, C and D genomes of oats (Linares et al., 1998; Ananiev et al., 2002; Liu et al., 2019), we could reveal the genome composition of polyploids and define the nature of major evolutionary changes in oat genomes. We can then discuss intergenomic translocations, their consequences for phylogeny, speciation and polyploidy, and creation of new hybrids and breeding.

### Re-evaluation of genome composition of tetraploid and hexaploid oat species and potential speciation in the genus *Avena*

Although the genome composition of hexaploid oat species is well known (Chen and Armstrong, 1994; Jellen et al., 1994; Liu et al., 2019), and also confirmed by our analysis using A, C and D genome-specific repetitive DNA probes, there are still debates about the classification and origin of genomes in tetraploid species. The genomic composition of tetraploid *A. magna, A. murphyi* and *A. insularis,* previously assigned as AACC (Leggett et al., 1994; Shelukhina et al., 2007), has recently been revised using molecular and cytological analysis (Peng et al., 2010; Yan et al., 2014; Tomaszewska and Kosina, 2021). Here, we presented new evidence confirming the CCDD genomic constitutions of the tetraploid *A. magna* and *A. murphyi* using published C genome-specific 45bp AF226603 probe (Ananiev et al., 2002; Liu et al., 2019) as well as D genome-specific Ast-T116 and Ast-R171 probes (Liu et al., 2019). These data together with high-density genetic markers analysis (Fominaya et al., 2021) suggested that the CCDD-genome tetraploids could contribute to the evolution of hexaploid oats (**Fig. 9**). Our *in situ* hybridization with pAs120 probe confirmed a clear dissimilarity of the two genomes in tetraploid *A. abyssinica* and *A. vaviloviana,* consistent with Irigoyen et al. (2001) and Yan et al. (2014), proving AABB genome composition of these species. Karyotype and molecular data revealed distinction of *A. agadiriana* from other species classified as AABB-genome tetraploids (Jellen and Gill, 1996; Badaeva et al., 2010; Peng et al., 2010; Yan et al., 2014; Chew et al., 2016). Our analysis showed evenly distributed signals of the D genome-specific Ast-T116 probe on 28 chromosomes of *A. agadiriana* suggesting that the genome composition of this species might be DDDD, not AABB as previously stated. On the other hand, pAs120 probe showed weak dispersed signals along all of the 28 chromosomes, thus, the evidence provided here should be taken as putative and encouraging for further exploration of this species. The phylogenetic analysis supported *A. agadiriana* being closely related to *A. canariensis* and *A. longiglumis* (Chew et al., 2016; Peng et al., 2018). Some authors speculated that the genome composition of this species might be AADD due to some similarities to CCDD and AACCDD *Avena* groups (Badaeva et al., 2010; Luo et al., 2018c), but our FISH signals of A and D genome-specific probes rather indicated an autopolyploid origin of *A. agadiriana*. Our results may also suggest that the genomes of *A. agadiriana* are different from those previously identified in other oat species as A, B, C and D and may require a different nomenclature (Yan et al., 2016), especially that the analysis of chloroplast genome demonstrated potential independent evolution of this species (**Fig. 9**; Fu, 2018). Correct recognition of the *A. agadiriana* genomes could complement the data on the origin of the D genome in AACCDD-hexaploid oats (Luo et al., 2018c), supported by easy crossing of these two species (Badaeva et al., 2010).

**Fig. 9.**
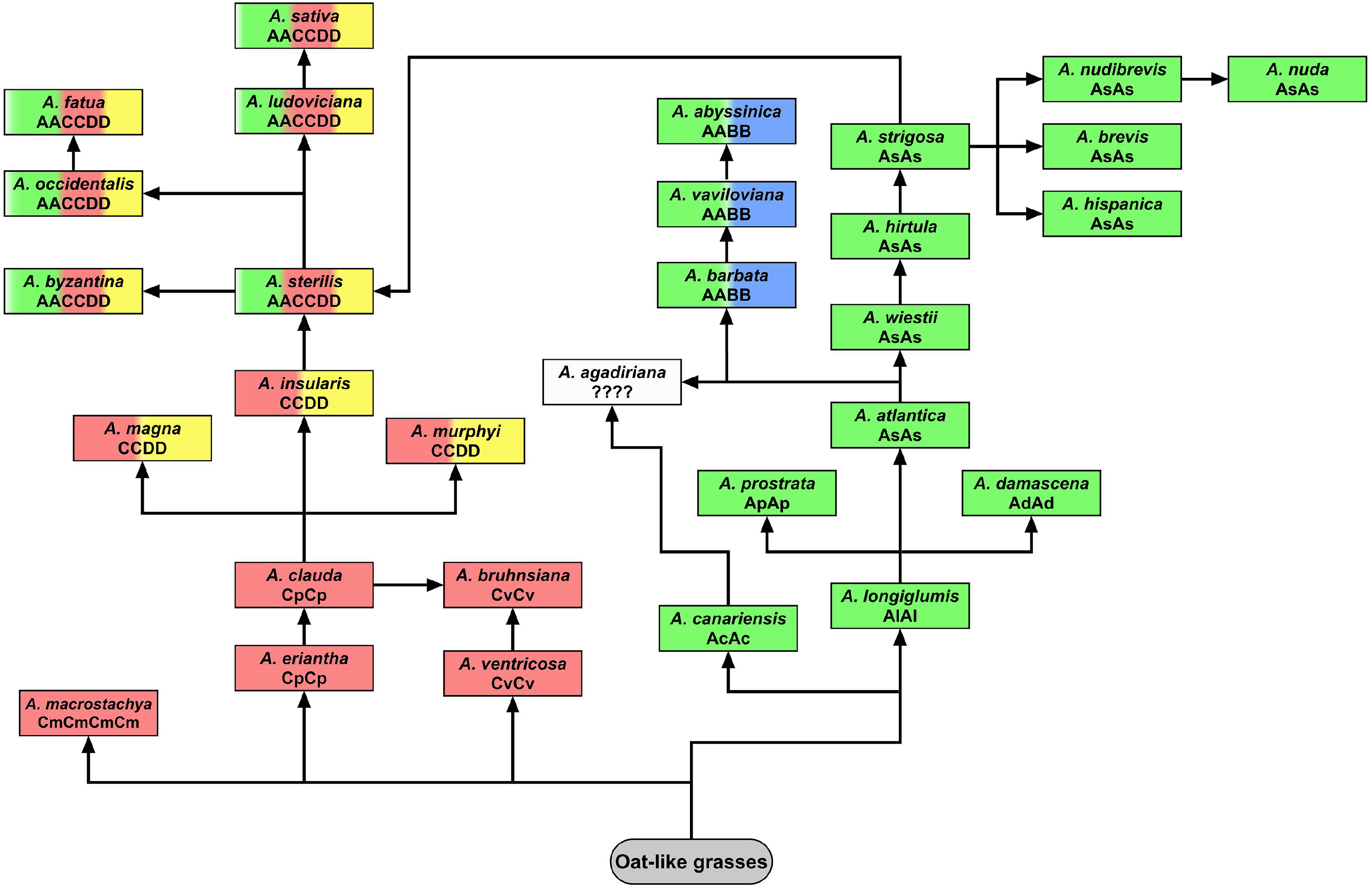
Model of potential evolutionary pathways leading to the development of modern tetraploid and hexaploid oat species based on the latest phylogenetic analyses. C and A genomes developed independently from oat-like grasses within the subgenus *Avenastrum* (Loskutov, 2007). According to Fu (2018), C-genome species diverged 19.9-21.2 million years ago (Mya), while *A. canariensis,* considered the oldest species having A genome, diverged 13-15 Mya. These data are contradictory to the analysis of chloroplast genome conducted by Liu et al. (2020), indicating that the diverged A genomes originate from *A. longiglumis. A. bruhnsiana* is an endemic species closely related to *A. ventricosa,* with *A. clauda* being a potential second ancestor (Gnutikov et al., 2022). Presumably different diploid species having the A and C genomes may have been involved in the evolution of polyploid oats (Peng et al., 2010). Species having AABB genomes (*A. barbata, A. vaviloviana, A. abyssinica*) show a close relationship with *A. wiestii* or *A. hirtula*. Peng et al. (2018) suggested that these species derived through autopolyploidization of As genome, contrary to Badaeva et al. (2010) and Irigoyen et al. (2001) who proposed allotetraploid origin. According to Peng et al. (2018), *A. agadiriana,* which developed independently, is closely related to the As and A_c_ genomes, but the genome composition of this species remains debatable (this paper; Badaeva et al., 2010; Yan et al., 2016; Luo et al., 2018c). Phylogenetic analysis conducted by Peng et al. (2018) indicated that C_p_ genome (*A. eriantha, A. clauda*) is more closely related to CCDD-tetraploids and AACCDD-hexaploids than Cv genome (*A. ventricosa*). Probably not one, but several of the extant CCDD-tetraploids were involved in the formation of hexaploid oat species (Yan et al., 2021), but *A. insularis* and diploid species having As genome seem to be putative ancestors of oat hexaploids (Drossou et al., 2004; Badaeva et al., 2005; Peng et al., 2018; Fominaya et al., 2021; Jiang et al., 2021). Molecular cytogenetic analysis revealed that none of the modern diploid oat species having A or C genome was the direct ancestor of polyploids, however, *A. insularis* showed a relatively high number of transcribed spacers, displaying sequence similarity to one accession of *A. hirtula* (Loskutov et al., 2021). On the other hand, the principal coordinates analysis revealed that *A. eriantha* and *A. longiglumis* are closer to CCDD oat tetraploids than other diploids (Yan et al., 2016). This was also proved by analysis of chloroplast and mitochondrial genomes (Fu, 2018), and easy crossing of *A. longiglumis* with CCDD tetraploids (see *A. magna* × *A. longiglumis* amphiploid in this paper; Drossou et al., 2004; Tomaszewska and Kosina, 2021).

### Identification of genomes in artificial hybrids and amphiploids of oats

Recognition of genome composition and genomic changes in hybrids is important in the context of plant speciation and evolution to explore drivers that played a major role in genome divergence (Alix et al., 2008; Patokar et al., 2016; Martin et al., 2020; Tomaszewska, 2021). The genome composition and intergenomic rearrangements of artificial hybrids and amphiploids of oats were investigated here for the first time using repetitive DNA sequences. A certain advantage of the repetitive DNA probes over the whole genomic DNA probes used for *in situ* hybridization is observed (Tomaszewska et al., 2022). Despite the often perfect discrimination of genomes in allopolyploids/amphiploids using total genomic DNA isolated from a diploid species (Tomaszewska and Kosina, 2021), such probes are not applicable to distinguish closely related genomes such as those observed in *Brassica* (Alix et al., 2008), *Triticum* (Tang et al., 2018), *Cenchrus* and *Urochloa* tropical forage grasses (Rathore et al., 2022; Tomaszewska et al., 2022) or highly homologous A and D genomes in *Avena* (Linares et al., 1998). The parental species of amphiploids studied here are meiotically compatible in the crossing process, exhibiting sometimes partial sterility (Loskutov, 2001), and these data together with established genome composition of hybrids and amphiploids contribute to a better understanding of the complex evolutionary processes within the genus *Avena*.

### Rapid genome changes in allopolyploids: intergenomic translocations in polyploid plant species and synthetic hybrids

Intra- and inter-genomic chromosome translocations have been recognized as significant processes accompanied natural evolution of diploid and polyploid plant species (Martin et al., 2020). The exchange of segments of non-homologous chromosomes occurred soon after allopolyploid formation, shape their genomes giving them an adaptive and evolutionary advantage (Tomaszewska, 2021). In some allopolyploid species, intergenomic translocations seem to be common, as in *Nicotiana* (Kenton et al., 1993), *Solanum tuberosum* and *S. caripense* (Braz et al., 2018), *Anthoxanthum* (Chumová et al., 2021), *Hordeum secalinum* (Bustos et al., 1996) and *H. capense* (Taketa et al., 1999) or wild wheats (Jiang and Gill, 1994; Badaeva et al., 1995), contrary to wheat cultivars (Badaeva et al., 2007) or *Secale* (Kubaláková et al., 2003) where intragenomic translocations located at pericentromeric regions of chromosomes, rarely interstitially, are the most common. In Musaceae, there are many translocations at any place in the chromosome arm (Wang et al., 2022).

Intergenomic translocations are a relatively common phenomenon in artificially induced allopolyploids (amphiploids). In newly synthesized wheat-rye allopolyploids, the transfer of rye telomeric chromatin involving the short arm of chromosome 1 (Kubaláková et al., 2003) to wheat centromere was observed in two continuous generations (Fu et al., 2010). Through distant crossings the yield of wheat cultivars has been improved and resistance to pathogens and insects has been induced. Numerous distal (terminal) intergenomic translocations were also detected in wheat-barley lines (Prieto et al., 2001; Nagy et al., 2002), *wheat-Aegilops biuncialis* amphiploids (Molnár et al., 2009), amphiploid trigeneric hybrid involving *Triticum, Thinopyrum* and *Lophopyrum* (Kosina and Heslop-Harrison, 1996) or in endosperm of *Avena* amphiploid (Tomaszewska and Kosina, 2021).

### Evolutionary dynamics in genus *Avena* in the context of intergenomic translocations

Comparative linkage mapping of oat species suggested extensive chromosome rearrangement in ancestral diploids, both involving SAT and non-SAT chromosomes (Badaeva et al., 2010; Latta et al., 2019); and intergenomic translocations occurred between non-homologous chromosomes were considered to be significant evolutionary forces leading to divergence of the tetra- and hexaploid oat species (Linares et al., 1998). Polyploid oat genomes are particularly rearranged showing multiple distal, rarely interstitial, intergenomic translocations. Our analysis of oat species and artificial hybrids having genomic composition of AACCDD showed that these two groups differ in the number and position of translocations, meaning that a new pattern of intergenomic translocations emerged soon after hybridization and was passed on and maintained over the next hybrid generations. The translocation pattern seen in amphiploids created by crossing different maternal diploids with paternal *A. sativa* indicated that some translocations were transferred from *A. sativa* to the hybrid generation. New translocations also appeared, supporting the hypothesis that at least some of the intergenomic translocations contributed significantly to the divergence of oat species. However, the analysis of *A. sativa* indicated ongoing genomic exchange in this hexaploid (Latta et al., 2019), thus, it is not only hybridization and genome duplication that affect genome rearrangement. In the hybrids involving *A. strigosa, A. eriantha,* and *A. magna,* a poor chromosome pairing was observed during meiosis, and the translocations may be the major factor limiting fertility rather than the lack of chromosome homology (Tomaszewska and Kosina, 2021). Thus, potentially, the presence of intergenomic translocations in *Avena* would limit backcrossing in breeding.

Our study showed that *A. abyssinica* and *A. vaviloviana* share some of the translocations between A and B genomes which indicates that translocations were one of the factors driving the divergence of AABB *Avena* group. The translocation of the entire arm of a single chromosome observed in *A. vaviloviana* may be species-specific or specific for this particular accession. Intergenomic translocations have not been identified in *A. agadiriana* using our selected probes, and the FISH pattern suggested that this species may be of autopolyploid origin thus its affiliation to the AABB group and potential drivers shaping this species remain debatable. Both our karyotype analysis and the data obtained by different authors (Jellen et al., 1994; Luo et al., 2018a, b; Tomaszewska and Kosina, 2021; Yan et al., 2021; Peng et al., 2022) indicated that some intergenomic translocations could also influence the divergence of the CCDD *Avena* group. Some translocations between the C and D genomes in hexaploid oat species involved the same chromosomes as in tetraploid CCDD species (this paper; Badaeva et al., 2010; Yan et al., 2021) supporting the hypothesis that one of the species belonging to this oat group contributed to the evolution of hexaploid *Avena* species. Unfortunately, without knowing the exact ancestors of hexaploids, it is difficult to verify whether translocations which appeared in CCDD *Avena* species were transferred to AACCDD species through hybridization. Analysis of intergenomic translocations across different hexaploid oat species and cultivars indicated that the pattern of intergenomic translocation of hexaploid *A. byzantina* is different from that observed in other wild, weedy or cultivated hexaploid oat species (Badaeva et al., 2011), and also ‘Kanota’ and ‘Ogle’ cultivars differed from each other in at least some of the intergenomic translocations (Jellen et al., 1994). Therefore, in oats, three types of intergenomic translocations should be distinguished: common or group-specific, species-specific, and cultivar- or accession-specific (Sanz et al., 2010).

### Intergenomic translocations and their implications for polyploid plant evolution

While some translocations are common for the Panicoideae group, *Setaria, Saccharum, Sorghum,* and *Zea* show further chromosome rearrangements which distinguish them from other grasses (Devos and Gale, 1997), and reciprocal translocations separating species within the *Secale* genus were revealed (Singh and Robbelen, 1977). Some of the intergenomic translocations observed in wheat group are species-specific, as those observed in *Triticum dicoccon, T. timopheevii* and *T. turgidum* (Jiang and Gill, 1994; Levy and Feldman, 2002), supporting the diphyletic hypothesis of the evolution of tetraploid wheats. Another examples of species-specific intergenomic translocations are those seen in tetraploid *Nicotiana tabacum* (Kenton et al., 1993). A nucleocytoplasmic interaction (NCI) hypothesis of polyploid plant genome evolution explains the presence of species-specific intergenomic translocations (Jiang and Gill, 1994). It was speculated that the newly formed polyploid has to overcome a bottleneck of hybrid sterility resulting from cytoplasmic-nuclear interactions of paternal and maternal species. Some chromosomal and/or genome changes must occur to stabilize genomes and restore fertility in hybrids. Intergenomic chromosome translocations seem to be one of the major mechanisms stabilizing non-homologous genomes that come together in polyploid nuclei.

### Intra- and inter-genomic translocations in plant breeding

The allopolyploid condition enables intra- and inter-genomic reshuffling of chromosomal segments and genes through translocations, recombination, transposition or introgressions (Kenton et al., 1993; Levy and Feldman, 2002; Patokar et al., 2016; Martin et al., 2021; Tomaszewska and Kosina, 2021; Tomaszewska et al., 2022). This unique feature, especially widespread in grasses, increases genome plasticity and enables to create new genetic combinations and genome arrangements. The fundamental research has implications for breeding oats if F1 hybrid combinations include undetected translocations leading to poor performance and restricting recombination; and in interspecific crosses, to exploit biodiversity through introgression from wild species, and perhaps enabling additional introgressions in wheat breeding. Agrawal et al. (2020) have developed pools of 20,000 oligonucleotides for use as probes in *Brassica*. By allowing identification of chromosome arms in chromosome preparations, these probes identified a reciprocal translocation present in one cultivar of *Brassica rapa,* cv. Purple Top Milan turnip, compared to cv. Chiifu-401 pak choi, which would make use of a hybrid difficult. It is likely that the translocations detected in the oat lines will mean care is required to use F1 hybrids in a breeding programme in case there are undetected translocations.

### Genome sequence assemblies

New long-molecule sequencing approaches are allowing assembly of *Avena* genome sequences, covering nearly 4,000Mbp in the diploids (Maughan et al., 2019; Liu et al. 2022; Peng et al., 2022). High coverage shows the reciprocal translocations which are also revealed with the chromosomal probes. Currently it is not possible to survey large numbers of lines, and too complex to reconstruct with high coverage of the recombinant chromosomes of multiple species (Fig. 9; chromosomal studies using *in situ* hybridization as Agrawal et al., 2020 or Huang et al., 2018 to identify translocations within species). It will be interesting to identify with base-pair resolution the chromosomal breakpoints, and further characterize the non-reciprocal nature of the translocations. We can then address key concern: what are the consequences of additional copies of alleles if segments are duplicated?

## Conclusions

Tetraploid and hexaploid oat species, as well as hexaploid and octoploid synthetic hybrids and amphiploids are characterized by multiple distal intergenomic translocations, in contrast to the wheat-group species (wheat, rye, barley) that show mainly intragenomic translocations located in centromeric and interstitial regions of chromosomes. The presence of distal, mostly non-reciprocal translocations in the *Avena* group makes oat chromosomes particularly rearranged. It suggests that intergenomic translocations were a major mechanism of divergence in the evolution of oat species, and a new pattern of translocations is established in synthetic hybrids and amphiploids. The consequences of distal and interstitial intergenomic translocations for hybrid stability and gene expression from both genomes involved in translocation remain little known.

## Acknowledgements

We are grateful to Bundesanstaltfür Züchtungsforschung an Kulturpflanzen (Braunschweig, Germany), National Small Grains Collection (Aberdeen, Idaho, USA) and Vavilov Institute of Plant Industry (St. Petersburg, Russia) for their generous provision of seeds.

## Funding

This project was funded under the RCUK-CIAT Newton-Caldas Initiative, with funding from UK’s Official Development Assistance Newton Fund awarded by UK Biotechnology and Biological Sciences Research Council (BB/R022828/1), and by the European Union’s Horizon 2020 research and innovation programme under the Marie Sklodowska-Curie grant agreement No 844564, individual fellowship to PT. This project has also received funding from the European Union’s Horizon 2020 research and innovation programme under the Marie Sklodowska-Curie grant agreement No 101006417.

## Notes

### Competing Interest Statement

The authors have declared no competing interest.

